# LOL: a Python package for leakage-aware phenotype prediction and confounding diagnostics in GEO transcriptomics datasets

**DOI:** 10.64898/2026.09.26.754596

**Authors:** Martin Nganga Muigano

## Abstract

Phenotype prediction from public gene-expression data is straightforward to implement but reliable prediction is harder. Leakage during feature selection can inflate performance estimates. Class imbalance can hide poor performance in minority classes. Confounds like tissue type, repeated sampling, and demographic differences can also create patterns that resemble phenotype-associated biology. Here, I present *LOL* (Latent Omics Learning), a Python package for phenotype prediction from bulk GEO RNA-seq and microarray data. LOL uses leakage-safe preprocessing, imbalance-aware evaluation, and group-aware validation where repeated samples are present. Each analysis produces two audit diagnostics. The leakage inflation index (ΔLII) measures the change in performance between naive and leakage-safe evaluation. PVCA-lite examines associations between the latent representation and available metadata covariates, including phenotype. I evaluated the workflow across six independent GEO cohorts. The datasets covered microarray and RNA-seq platforms, binary and multi-class phenotypes, and sample sizes ranging from 18 to 566. The analyses produced a varied set of outcomes. Across diverse GEO datasets, LOL can reveal phenotype-associated expression patterns while identifying factors distort their interpretation. These include information leakage, class imbalance, repeated sampling, tissue differences, and demographic or other metadata-associated confounding. The resulting diagnostics help distinguish robust predictions from results that require further scrutiny. LOL does not introduce a new representation-learning algorithm. Its contribution is a reproducible workflow that places validation and diagnostic checks alongside prediction. The resulting analysis can expose weaknesses in an apparently convincing result and make those limitations visible. LOL is freely available under the MIT license from https://github.com/ngangao/lol-omics.

## 1. Introduction

Latent representation learning methods such as principal component analysis (PCA) and variational autoencoders (VAEs) are useful in compressing high-dimensional gene-expression matrices prior to downstream analysis (Teschendorff et al., 2007; Yamamoto et al., 2009). However, when they are applied to gene expression public repositories like GEO, standard machine learning workflows frequently fall into a hidden trap. The models might achieve inflated cross-validation performance by accidentally memorizing source identity, laboratory processing runs, or technical batch structures rather than true underlying biology (Parker and Leek, 2012; Soneson et al., 2014). So, without rigorous variance auditing and leakage-safe cross-validation, published biomarker signatures may risk representing methodological artifacts rather than robust biological insights.

In transcriptomic phenotype prediction, three methodological oversights are easily neglected that are likely to compromise the results. The first one involves fitting feature selection and latent dimensionality reduction globally across the entire dataset prior to cross-validation. This might introduce data leakage, improper data splitting, and inflated model performance (Smialowski et al., 2010; Starcke et al., 2025). Second, relying solely on overall accuracy or standard ROC-AUC on imbalanced cohorts risks masking a total failure to predict minority classes, especially for imbalanced data (Jeni et al., 2013; Imani et al., 2026). Lastly, there is often the risk of interpreting high classification scores as biological signals without auditing for underlying confounds like sequencing platforms (Soneson et al., 2014; Chi et al., 2026). Other common confounding features include batch artifacts, laboratory processing runs, or repeated subject sampling. To resolve these vulnerabilities, I introduce LOL, a Python framework for leakage-aware phenotype prediction and confounding diagnostics on GEO transcriptomics datasets. The primary contribution of LOL is not in algorithmic novelty, but in enforcing rigorous pipeline hygiene.

The LOL framework addresses the challenges highlighted above through a three-stage architectural pipeline. First, it implements diagnostic Principal Variance Component Analysis (PVCA-lite) to explicitly quantify latent variance drivers across high-dimensional feature spaces. This allows a priori determination of whether predictive signals are dominated by biological phenotypes, technical batch artifacts, or patient identity leakage. Second, LOL replaces naive sample-level splitting with group-aware, fold-safe cross-validation structures that strictly isolate repeated measures and multi-biopsy donor dependencies. Finally, it provides automated feature loading extraction and visualization tools to evaluate whether the underlying latent space preserves coherent biological signatures. LOL’s central contribution is its two diagnostics that make the risk visible as an initial, automatically computed output: a leakage inflation index (ΔLII) and a PVCA-lite confound diagnostic.

## 2. Methodology

### 2.1. Pipeline implementation

LOL accepts a GEO accession or a local expression matrix together with a phenotype label for each sample. The pipeline has four steps. The first step is normalization. Here, RNA-seq counts are converted to log-CPM. For gene i and sample j, raw count x□□ is converted as follows.

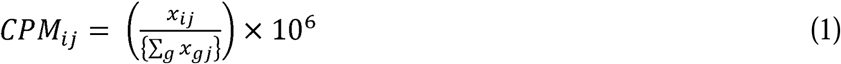

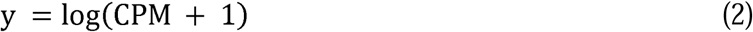

Where *CPM_ij_* is the normalized expression value in counts per million for gene *i* in sample *j*. ∑*_g_x_gi_* is the sum of raw counts across every gene *g* in sample *j* while *y* is the final value that LOL uses downstream after normalization.

Microarray intensities are then normalized by quantile normalization instead. Platform is detected automatically from whether values are non-negative integers. In step two, highly-variable-gene selection is performed. LOL keeps the N genes with the highest variance across samples. This step controls memory use and runtime. Step three involves latent extraction. PCA is the default method for this. Still, a small VAE is available as an alternative. The VAE is trained by minimizing the following loss.

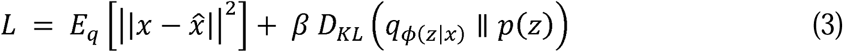

Where,

- L = total quantity minimized while training the VAE
- *x* = one sample’s original expression vector (input to the encoder)
- *x̂* = the decoder’s reconstruction, i.e. what the VAE predicts looked like after compressing it to ( and decompressing back
- *x* - *x̂* = squared reconstruction error between the original and the reconstruction
- *z* : the latent (compressed) representation of the sample
- the encoder’s distribution over *z* given input *x* with weights
- *D_KL_* = the Kullback-Leibler divergence, a measure of how far the encoder’s distribution is from that prior
- β: a tuning constant controlling how strongly the model is pushed to match the prior versus reconstructing accurately

Given that the term *D_KL_* is the Kullback-Leibler divergence between the approximate posterior and a standard normal prior, for a Gaussian posterior with mean μ and variance σ², this divergence has a closed form.

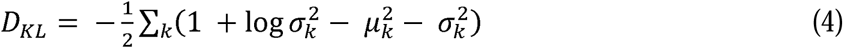

Where *k* is the index running over each latent dimension *μ_k_* is the mean that the encoder outputs for latent dimension, and 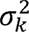 is the variance the encoder outputs for latent dimension. Latent samples are drawn with the reparameterization trick, z = μ + σ ⊙ ε, where ε is drawn from a standard normal distribution. PCA and the VAE are offered side by side. PCA is useful in reducing non-linear dimensionality (Tsuyuzaki et al., 2020) while VAE help to incorporate latent variables (Gomari et al., 2022). Step four is classification. A Random Forest classifier is trained on the latent features. In this step, class weights are balanced by default.

Evaluation is nested by default. Gene selection and the latent fit are repeated inside each cross-validation fold, instead of once on the full dataset before splitting. Reported metrics include macro-F1 alongside accuracy and ROC-AUC. Macro-F1 is less sensitive to class imbalance than accuracy. When repeated samples from one subject are present, an optional group identifier routes cross-validation through scikit-learn’s StratifiedGroupKFold. Every sample from one subject then stays on the same side of each fold.

Two data-loading paths are supported. Most microarray series embed a processed data table directly in the series matrix. LOL reads this table with load_series_matrix. Files are fetched with geofetch (Khoroshevskyi et al., 2023) but manual download can be performed. Many RNA-seq series submit processed data as separate supplementary files, typically a genes-by-samples counts table and a separate sample-metadata file. load_counts_table handles this shape. Both loading paths were run against real, previously untested code during development of the validation set in Section 3. This surfaced and fixed two geofetch command-line errors and it motivated the addition of load_counts_table.

### 2.2. Permutation testing

A cross-validated score alone does not show that a phenotype is distinguishable from chance. LOL reruns the full nested pipeline on randomly shuffled labels, 30 by default, and reports where the real score falls relative to the resulting null distribution. The estimated p-value is as follows.

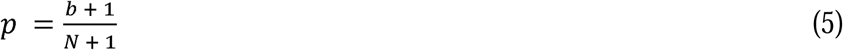

Here, b is the number of null scores at or above the real score, and N is the number of permutations. When a subject-grouping identifier is supplied, shuffling is done at the subject level. Each subject’s one true label is shuffled and that shuffled label is propagated to all of that subject’s samples. A subject’s diagnosis is one fact about them, not an independent draw per sample. So, this step requires each group to carry one consistent label. LOL raises an explicit error when it does not, rather than returning a misleading result (Section 3.6).

### 2.3. Leakage inflation index (ΔLII)

For a given metric M (accuracy, macro-F1, or ROC-AUC), the leakage inflation index is defined as follows.

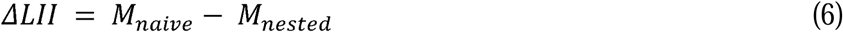

M_naive is the score without nested refitting. M_nested is the score from the leakage-safe evaluation. LOL computes both from the same fitted pipeline and reports the difference directly. A value near zero means the naive shortcut did not change the result on this dataset. A large positive value means it did, and the naive score should not be reported. Both outcomes are informative. Section 3 reports examples of each.

### 2.4. PVCA-lite confound diagnostic

Principal Variance Component Analysis (PVCA) partitions the variance of each principal component among a set of covariates. It uses a mixed-effects model fit jointly across all covariates. PVCA has been used to screen microarray and RNA-seq data for batch effects (Li et al. 2009). A full joint implementation needs a mixed-model fitting library, a heavier dependency than the rest of LOL requires. In this study, I implemented a simplified, per-covariate approximation instead.

For each covariate c and each latent dimension j, LOL computes an R² value. For a categorical covariate, this is a one-way-ANOVA R².

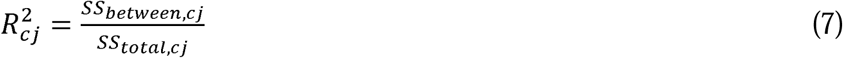

Where;

- 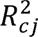 = fraction of variance in latent dimension *j* explained by covariate
- *SS_between,cj_* = how much the group means (one mean per category of c, e.g. per tissue type) differ from the overall mean
- *SS_total,cj_* = how much every individual sample’s value differs from the overall mean, i.e. the total variance available to explain

For a continuous covariate, R² is the squared Pearson correlation between the covariate and the latent dimension.

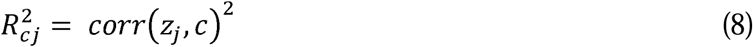

Where *z_j_* is the values of latent dimension across all samples, *c* the values of the covariate across all samples (e.g. age in years), and *corr*(*z_j_*, *c*) is the Pearson correlation coefficient between them. LOL then computes a weighted average across latent dimensions. The weight for dimension j is its share of total explained variance.

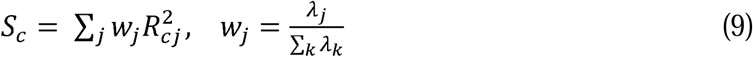

Here, λ_j is the explained variance of principal component *j*, and S_c is the reported share for covariate *c*. Covariates are handled independently, not jointly. Their shares are not adjusted for overlap with each other, and shares are not constrained to sum to one. Two correlated covariates can each show a substantial, overlapping share.

## 3. Validation

The LOL pipeline was evaluated using six independent case studies. Each comprises real GEO datasets and illustrates different capabilities of the tool. All six results include a permutation test comparing the real score to a null built from the full nested pipeline run on shuffled labels. Across the six GEO cohorts, nested classification performance varied across datasets and phenotype definitions (Table 1). Macro-F1 ranged from 0.642 to 1.000. Permutation tests reached significance in four cohorts. GSE22619 did not show evidence of above-chance classification while the remaining datasets showed significant classification performance (Figure 1). GSE66407 was found to have a confound and therefore a pooled p-value is reported.

**Figure 1:**
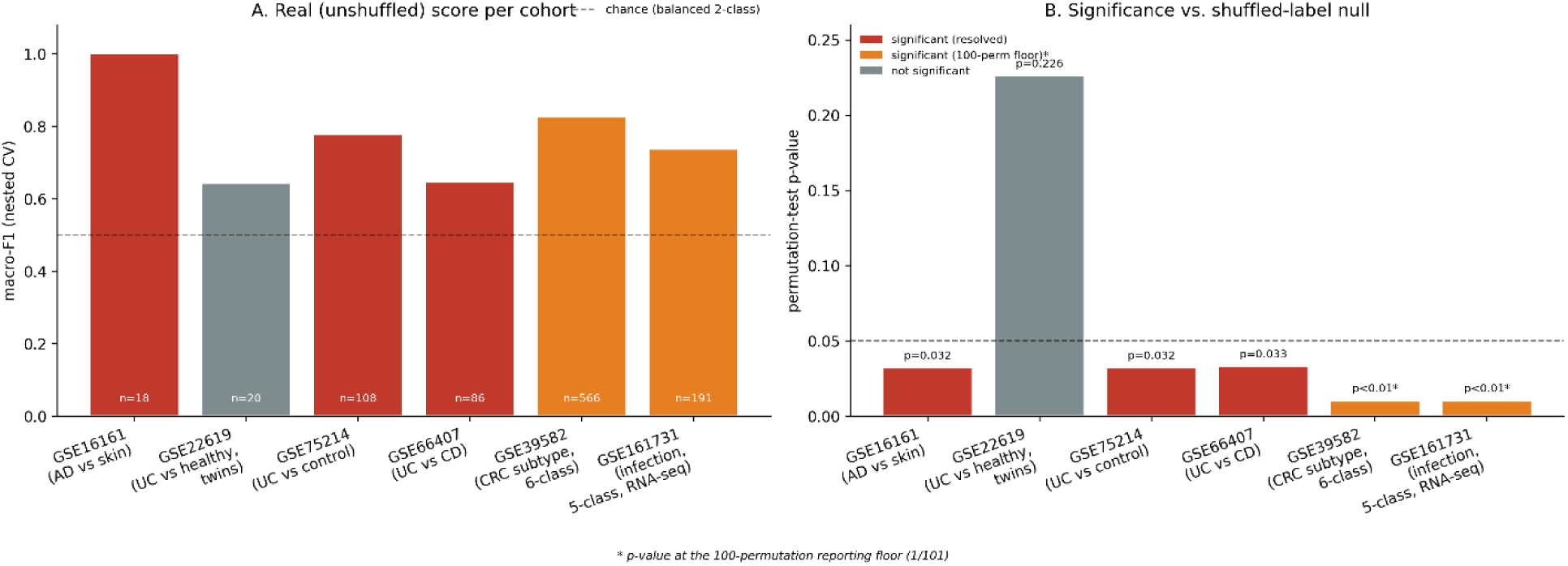
Validation summary across six real GEO cohorts. (A) Real, unshuffled, nested-CV macro-F1 per cohort. (B) Permutation-test p-value per cohort.

**Table 1.** Datasets used in the validation alongside the core results per cohort.

| Dataset | Comparison | n (retained after QC) | macro-F1 | p-value |
| --- | --- | --- | --- | --- |
| GSE16161 | AD vs. healthy skin | 18 | 1.000 | 0.032 |
| GSE22619 | UC vs. healthy, discordant twins | 20 | 0.642 | 0.226 |
| GSE75214 | UC vs. control, colon tissue | 108 | 0.777 | 0.032 |
| GSE66407 | UC vs. Crohn's, matched colon tissue | 86 | 0.646 | 0.033 (pooled) |
| GSE39582 | Colorectal cancer subtype (6-class) | 566 | 0.826 | <0.01 |
| GSE161731 | Infection cohort (5-class, RNA-seq) | 191 | 0.737 | <0.01 |

The diagnostic results also showed substantial variation in leakage inflation and the metadata factors associated with the latent representations (Table 2). ΔLII was positive in four cohorts. This points to higher macro-F1 under the naive evaluation than under the leakage-safe procedure. The largest inflation occurred in GSE66407 while GSE22619 showed a small negative ΔLII. PVCA-lite identified phenotype as the leading covariate in GSE16161, GSE75214, and GSE39582. However, other datasets showed stronger associations with non-phenotypic covariates. Twin-pair identity dominated the latent structure in GSE22619 and inflammation was the leading covariate in GSE66407. For GSE161731, the cohort, race, and ag contributed substantially to the latent structure. Tissue was constant in GSE16161 and GSE75214 and therefore could not be assessed as a source of between-sample variation. These results show that classification performance was accompanied by different patterns of latent-space structure across the six cohorts.

**Table 2.**
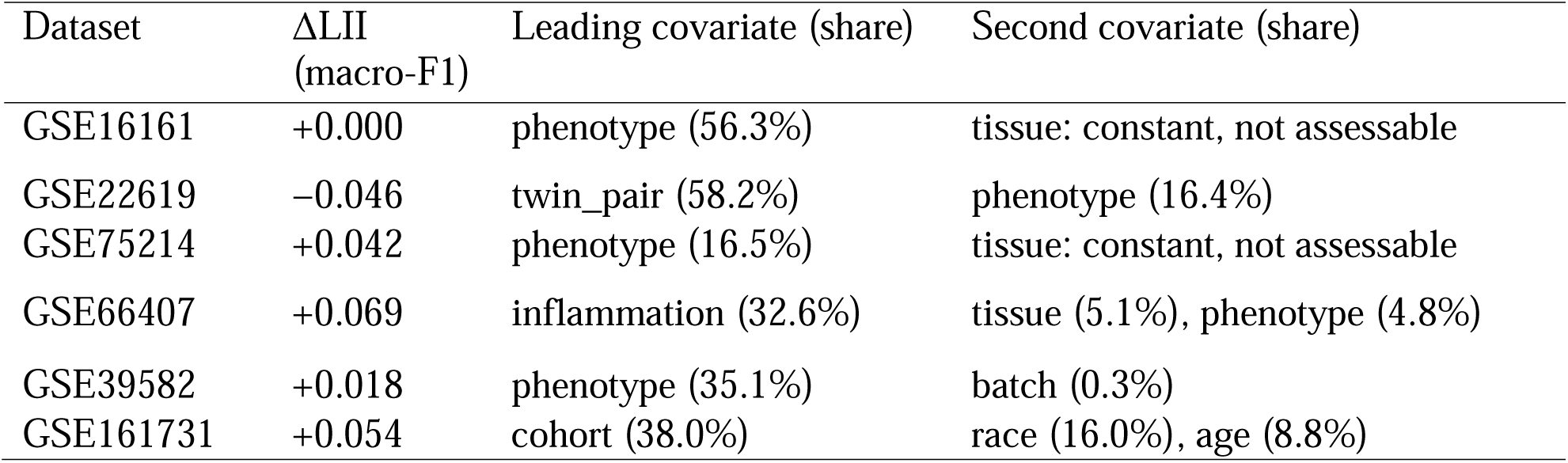
ΔLII (macro-F1) and PVCA-lite results for all six cohorts.

### 3.1. Case study 1: GSE16161, atopic dermatitis versus healthy skin

GSE16161 is a microarray series. It comprises 18 skin biopsies of 9 samples from patients with atopic dermatitis and 9 from healthy controls. Labels were built from the series matrix characteristics field. Initial phenotypic scrutiny produced a perfect 1.000 accuracy, ROC-AUC, and macro-F1. These results called for further scrutiny on all 18 samples. Refitting inside each fold did not change this result. In the case of ΔLII, the values were 0.000 for all three metrics (Table 2). A 30-permutation test confirmed the score was not due to chance because every shuffle scored below the real result (p = 0.032).

Latent representation showed a clear phenotype-associated structure. PVCA-lite indicated that phenotype accounted for 56.3% of the variance in the latent space. Tissue was the constant covariate across all samples and therefore could not account for the separation. To examine the features underlying this separation, I extracted the strongest positive and negative PC1 loadings from the fitted pipeline (Figure 2). The corresponding expression patterns showed opposing signatures across the two phenotype groups with the same features that contributed most strongly to PC1 displaying distinct expression patterns between samples. This provides a direct view of the molecular structure underlying the classification result rather than relying only on the performance metrics. The absence of measurable leakage inflation, the permutation result, the phenotype-associated latent structure, and the expression patterns of the leading PC1 features support a strong transcriptomic distinction between atopic dermatitis and healthy skin. This likely explains the perfect recall and accuracy.

**Figure 2.**
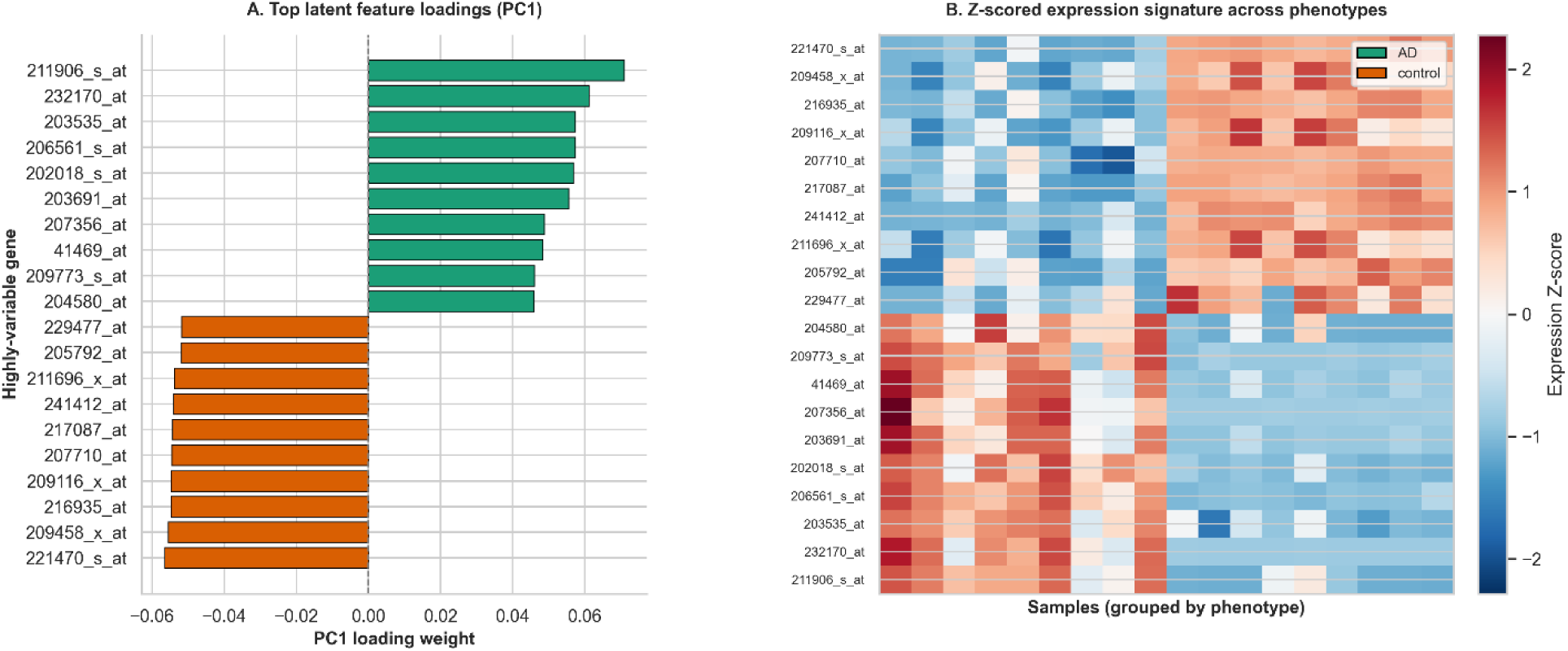
Latent feature structure underlying phenotype prediction in GSE16161. (A) The ten most positive and ten most negative PC1 loading weights among the highly variable features used by LOL. (B) Row-wise Z-scored expression of these PC1 driver features across the 18 samples, ordered by phenotype.

### 3.2. Case study 2: GSE22619, ulcerative colitis in discordant twins

GSE22619 is an Affymetrix microarray series of 20 samples from 10 discordant twin pairs: one healthy twin and one twin with UC per pair. Labels were built from sample titles. ΔLII showed a mixed pattern by metric whereby the accuracy and macro-F1 were both slightly higher nested than naive. The ROC-AUC was substantially higher naive than nested (0.810 versus 0.665, ΔLII = +0.145). Reporting only accuracy would have missed this. A 30-permutation test gave a clean null: macro-F1 0.642, p = 0.226. This was above the 0.05 threshold expected for twins matched on genetic background. PVCA-lite gave independent, structural evidence for the same conclusion. Twin-pair identity accounted for 58.2% of latent-space variance, which was more than three times the 16.4% accounted for by UC-versus-healthy status. For this cohort, the family from which a sample comes from explained more of the latent representation than disease status does. The classification null and the confound finding corroborate each other.

### 3.3. Case study 3: GSE75214, ulcerative colitis versus control in colon tissue

GSE75214 is also an Affymetrix microarray series. The dataset contains three diagnosis groups namely control, UC, and Crohn’s disease. These are distributed across colon and ileum tissue types. An initial analysis pooled colon and ileum control samples against colon-only UC cases. This produced a tissue-type confound since colon and ileum expression differ substantially regardless of disease. So, the analysis was restricted to colon tissue only, and to UC versus control. This left 108 samples (97 UC and 11 control). Restricting to matched colon tissue changed accuracy slightly from 0.944 naive to 0.935 nested (ΔLII = +0.009). This showed that the tissue confound was not the primary driver of this result. Further analysis showed that the Macro-F1 was 0.777, which was below the accuracy (0.935) and ROC-AUC (0.993) metrics. Per-class recall explains this observation in that only 45% of true controls correctly identified. Eleven samples in the minority class were below the sample size at which classifier performance was stable for this kind of task. A permutation test confirmed result (p = 0.032). PVCA-lite gave phenotype a 16.5%. This points to an imbalanced, limited signal with tissue being the only constant in this colon-only subset. The influence of latent feature dimensionality and cohort size on classification performance was also investigated for this cohort. I first compared the predictive efficacy of PCA and VAE compressions across multiple latent dimensions against a baseline of 2000 raw highly variable genes. Subsequently, I performed a subsampling analysis to assess the model’s sensitivity to reduced sample sizes while quantifying the degree of performance inflation (ΔLII) caused by data leakage during cross-validation. The resulting performance trajectories and leakage evaluations are presented in Figure 3.

**Figure 3.**
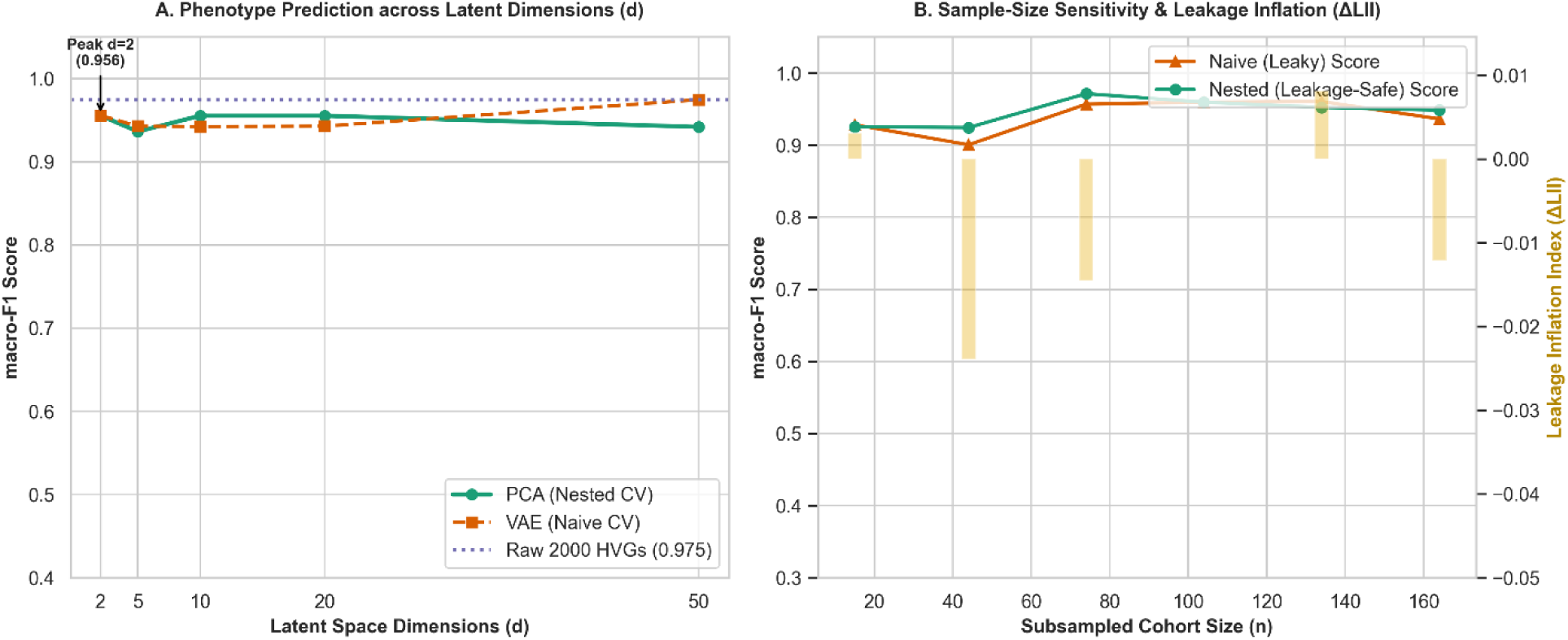
Impact of latent dimensionality and sample size on phenotype prediction and data leakage. (A) Macro-F1 scores for classifying inflammatory bowel disease phenotypes (Ulcerative Colitis vs. Crohn’s Disease) across varied latent space dimensions (d). Performance compares PCA with nested cross-validation (CV) against a Variational Autoencoder (VAE) with naive CV, benchmarked against a baseline of 2000 raw Highly Variable Genes (HVGs). (B) The sensitivity of macro-F1 scores to subsampled cohort sizes (n), contrasting naive (leaky) and nested (leakage-safe) CV pipelines.

### 3.4. Case study 4: GSE66407, ulcerative colitis versus Crohn’s disease

GSE66407 is an Affymetrix microarray series comprising 368 gut biopsies from 109 patients. It included patients with UC, Crohn’s disease, and control diagnoses across multiple colon and ileal segments. UC samples cluster in rectum and sigmoid colon while Crohn’s samples are disproportionately ileal. This is consistent with each disease’s known pathophysiology. Yet, this could be a source of a confound for classification. The analysis in this case was restricted to three tissue-balanced mid-colon segments (ascending, transverse, descending colon), which gave 86 samples (48 UC and 38 Crohn’s disease). Patients in this cohort contribute up to nine biopsies each. Cross-validation and permutation test were run with patient identity as the grouping variable. This helped to avoid patient’s biopsies being split across either a fold or a shuffle. In the pooled result, nested macro-F1 was 0.646 while ΔLII was +0.069. However, there was a real gap between naive (0.715) and nested (0.646) evaluation. A 150-permutation test gave p = 0.033. Applying the same PVCA-lite check used on previous cohorts surfaced something the original tissue- and patient-matching had not addressed: inflammation status. It accounted for 32.6% of latent-space variance, way above both tissue (5.1%) and phenotype (4.8%). A cross-tabulation showed that inflammation status was unevenly distributed between diagnoses even within this matched tissue subset (UC: 12 inflamed, 35 non-inflamed; Crohn’s: 17 inflamed, 21 non-inflamed) as shown in Figure 4. This is possibly because Crohn’s disease is segmental while UC inflammation is continuous. Therefore, a given mid-colon segment can be inflamed in UC more often than in Crohn’s even within matched tissue. The classification was rerun separately within each inflammation stratum. Neither reached significance: inflamed-only (n = 29, macro-F1 0.561, p = 0.161) and non-inflamed-only (n = 56, macro-F1 0.562, p = 0.194). The lack of conclusive results means that one cannot conclude that the pooled significance was substantially inflammation-driven. So, GSE66407’s result presents as inconclusive case. A larger inflammation-matched cohort could help to resolve this. Nevertheless, the case study argues against relying on a single diagnosis due to the risk of mechanically driving the result in one direction.

**Figure 4.**
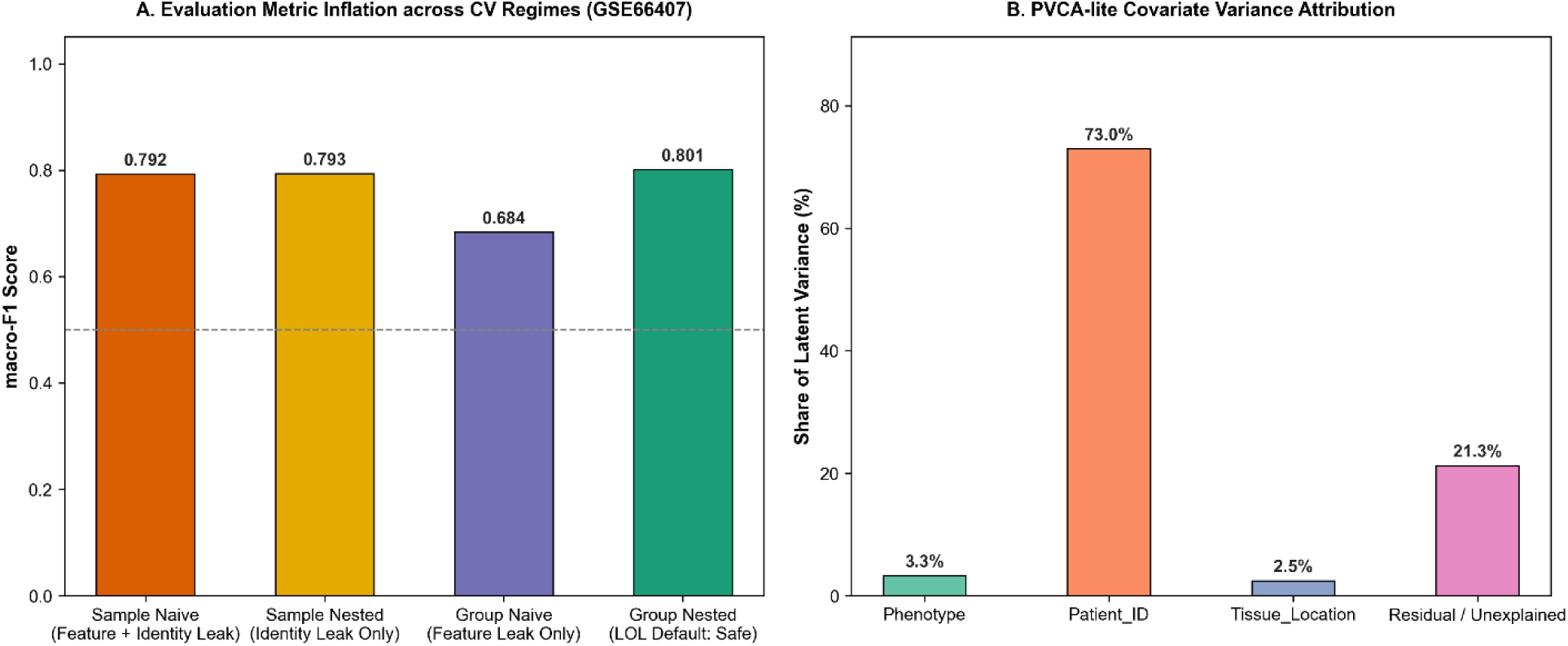
A leakage audit on GSE66407. (A) Evaluation Metric Inflation across CV Regimes. (B) PVCA-lite Covariate Variance Attribution.

### 3.5. Case study 5: GSE39582, colorectal cancer molecular subtype

The next case involves an Affymetrix microarray series with 585 samples. After removing 19 non-tumoral controls, a dataset of 566 primary colorectal tumors was retained. This was the largest and only 6-class cohort in this study. The classification was based on Cartes d’Identité Tumeurs (CIT) molecular subtyping system of transcriptomic classification framework (Marisa et al., 2013). The system categorizes colon cancer into six molecular subtypes (C1 to C6). Applying the LOL framework for this dataset produced a nested macro-F1 of 0.826 and a ΔLII of +0.018 (naive 0.844, nested 0.826). Naive and nested scores agreed closely at this sample size. A 100-permutation test confirmed the result was not due to chance (p < 0.01) with all 100 null shuffles falling below 0.174. PVCA-lite was run against real clinical covariates (e.g. tumor location, discovery-versus-validation batch, sex, MMR status). Phenotype accounted for 35.1% of variance association while batch accounted for only 0.3%. The phenotype was ahead of every technical covariate in the study. However, one caveat applies here. The C1-C6 labels were originally derived by unsupervised clustering on this dataset’s own expression data. This makes the task closer to recovering a clustering structure from similar data than to predicting an independently-diagnosed clinical phenotype.

### 3.6. Case study 6: GSE161731, whole-blood RNA-seq infection cohort

Lastly, LOL was used to evaluate phenotypes in GSE161731. This was the only RNA-seq cohort in this study. The dataset comprised the transcriptome of the peripheral blood of 46 subjects with COVID-19 infection. The GEO dataset had 198 samples of which 3 had failed QC. Another two were removed during the analysis in this study. So, a final dataset of 191 samples was analyzed from the infection cohort (COVID-19, seasonal coronavirus, bacterial pneumonia, influenza, or healthy). The analysis of this RNA-seq data gave a nested macro-F1 of 0.737 with ΔLII = +0.054 (naive 0.791, nested 0.737). A 100-permutation test confirmed that the result was not due to chance (p < 0.01). All 100 null shuffles fell below 0.215. After excluding the two conflicting subjects, PVCA-lite against real demographic covariates showed that the cohort accounted for 38.0% of variance association, race for 16.0%, and age for 8.8% (Figure 5). Before that exclusion, age was the single largest source of latent-space variance at 49.2%, ahead of cohort itself at 37.6%. Checking age by cohort directly explains why: healthy controls average 18.4 years old with a standard deviation 0.6. So, most were aged 18 while bacterial pneumonia patients averaged 61. So, the strong, statistically significant result cannot be attributed exclusively to infection-type signal alone.

**Figure 5.**
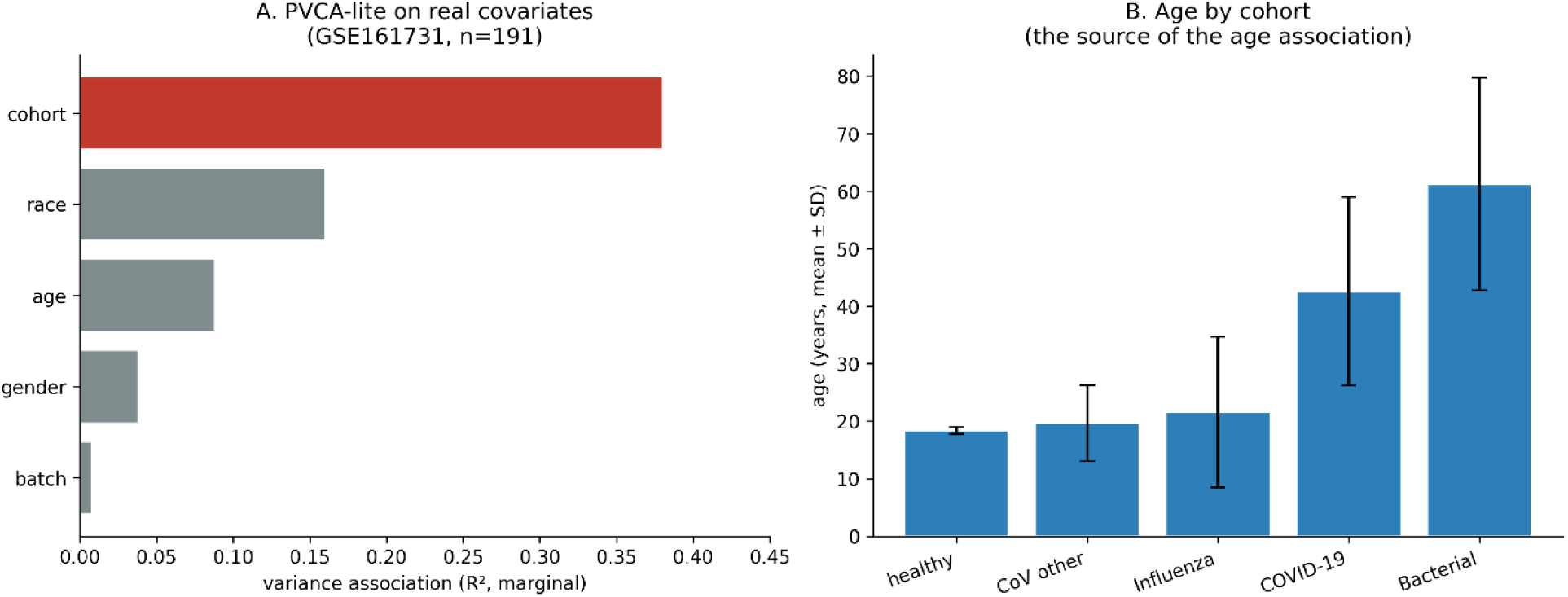
Confound diagnostic on GSE161731. (A) PVCA-lite variance association per covariate after label cleanup. (B) Mean age by cohort, error bars show standard deviation.

### 3.7. Reproducibility across cohorts

GSE16161, GSE22619, and GSE75214 were each analysed twice. The first analysis used purpose-built script during initial development. The second used the package’s standard Pipeline.evaluate() and Pipeline.permutation_test() methods. The ΔLII and PVCA-lite figures reported above were computed in a third pass once those diagnostics existed. GSE75214’s core evaluation result reproduced exactly across passes. GSE16161 and GSE22619 reached the same conclusion in every pass: p = 0.032 each time for GSE16161, and a clean null each time for GSE22619 (p = 0.16, then p = 0.226).

## 4. Discussion

This work involved the development of a lightweight pipeline for bulk GEO phenotype prediction. The goal of this pipeline is to capture confounding factors prior to downstream analysis. Various case studies presented show that predictive performance alone is not enough to interpret phenotype classification from public transcriptomic data. Gene expression datasets contain multiple sources of biological and technical variation, collectively known as confounding factors (Bruning et al., 2016; Toubia et al., 2018; Hatano et al., 2025). Some of this information may or may not be related to the phenotype of interest. For example, age, tissue type, inflammation, patient characteristics, experimental batch, and other study-specific factors can influence gene expression patterns. If these factors are associated with the outcome, a classifier can learn them as if they were biological signals of interest. This problem has long been recognized in transcriptomic studies (Leek et al., 2010; Goh et al., 2017; Papiez et al., 2019; Cote et al., 2022; Yu et al., 2024). Such unmodelled sources of variation can produce substantial changes in gene expression patterns and affect biological conclusions.

Heterogeneity is particularly important when working with GEO datasets. GEO provides access to a large and heterogeneous collection of publicly available expression studies (Edgar et al., 2002). While this makes it valuable for secondary analysis, it also means that one has to consider the risk of substantial variation in the data for cross-cohort comparison. The heterogeneity comes in many forms such as study design, sample composition, tissue types, clinical characteristics, and experimental procedures. Thus, variables that were incidental to the original research question may become important predictors in a secondary classification analysis. Previous work has shown that non-biological variation like technical variables can strongly influence genomic prediction when correlated with a variable of interest (Goh et al., 2017).

GSE66407 and GSE161731 illustrate a related but distinct problem. For this the GSE161731 dataset, the study objective was to distinguish immune response to COVID-19 infection for individuals categorized as having seasonal coronavirus, influenza, bacterial pneumonia, or healthy controls. The classification signal was strong but the clinical groups differed substantially in age. So, further scrutiny was needed. Since age was associated with extensive transcriptional variation, a model trained to on this dataset may capture age-related expression patterns instead of infection-related patterns. In the case of GSE66407, separation between Crohn’s disease, ulcerative colitis, and control was found to be strongly affected by tissue sources and inflammation status. The GSE66407 analysis highlights the value of examining potential confounders systematically. Confounding of this type has been recognized in gene expression literature where age and other sample characteristics are correlated with disease status (Leek et al., 2010; Hatano et al., 2025). LOL captures confounding factors through a prediction that reports ΔLII or a PVCA-style confound diagnostic as a built-in output. The role of the diagnostic component in LOL is not to remove every source of variation automatically. Instead, it provides a simple way to determine whether available metadata variables are associated with the major structure of the expression data before biological conclusions are drawn.

This study has several limitations worth highlighting. First, even though LOL standardizes the main analytical workflow, it is not a fully automated one-size-fits-all solution. Instead, dataset-specific manual curation is still needed to interpret metadata, resolve ambiguous labels, identify problematic samples, and determine appropriate covariates or grouping variables. This traces back to the nature of the GEO datasets as being heterogeneous in nature. Thus, the reproducibility provided by LOL applies primarily to the analytical workflow while core aspects of data preparation remain dependent on user judgment. Second, PVCA-lite is a marginal approximation as opposed to a joint variance decomposition. This means that contributions from correlated covariates may overlap. Third, some datasets contained relatively few samples, which might limit the ability to distinguish weak biological signals from confounding factors. Fourth, the current implementation supports out-of-sample transformation and nested evaluation for the PCA-based latent representation. However, the VAE pathway does not yet provide an equivalent out-of-sample transformation. Fifth, LOL is currently designed for bulk microarray and RNA-seq data and does not address single-cell or single-nucleus datasets. Finally, labels derived from expression data can introduce circularity when the same molecular information is subsequently used for prediction. Such results should therefore be interpreted as associations with the derived phenotype rather than independent validation of it.

## 5. Conclusion

LOL provides a small, documented and leakage-safe pipeline for testing whether phenotypes can be predicted from bulk GEO gene-expression data. It includes two diagnostic measures for assessing potential leakage and confounding. These checks were applied consistently across independent datasets. Across six independent cohorts covering microarray and RNA-seq data, binary and multi-class phenotypes, and a wide range of sample sizes, LOL identified several different outcomes. These included a leakage-driven inflation of performance, a negative result supported by independent checks, and biological and demographic confounders. This variation is important because it shows that applying the same workflow does not necessarily produce uniformly positive results. Instead, it can reveal when an apparently convincing prediction requires further scrutiny or cannot be interpreted with confidence. LOL is not intended to replace existing methods for latent representation learning. Its main value lies in combining prediction with systematic checks of how that prediction was obtained and what factors may be contributing to it. In this sense, LOL provides a practical framework for asking whether a phenotype can be predicted and how confidently that prediction can be interpreted.

## Data Availability

All gene-expression datasets analysed in this study are publicly available from the NCBI Gene Expression Omnibus (GEO) https://www.ncbi.nlm.nih.gov/geo/ under the following accession numbers: GSE16161, GSE22619, GSE75214, GSE66407, GSE39582, and GSE161731. Each accession page includes the original submitters’ sample-level metadata used to build phenotype labels and covariates in this study. The LOL source code is available under the MIT license at https://github.com/ngangao/lol-omics.

## References

Bruning, O., Rodenburg, W., Wackers, P. F. K., van Oostrom, C., Jonker, M. J., Dekker, R. J., et al. (2016). Confounding Factors in the Transcriptome Analysis of an In-Vivo Exposure Experiment. PLoS One 11, e0145252. doi: 10.1371/journal.pone.0145252

Chi, H., Liu, X., Yan, Y., Wang, L., Luo, P., Su, L., et al. (2026). Trustworthy AI for disease pathogenesis and precision medicine: A disease-agnostic framework from data to deployment. Genes & Diseases, 102480. doi: 10.1016/j.gendis.2026.102480

Cote, A. C., Young, H. E., and Huckins, L. M. (2022). Comparison of confound adjustment methods in the construction of gene co-expression networks. Genome Biol 23, 44. doi: 10.1186/s13059-022-02606-0

Edgar, R., Domrachev, M., and Lash, A. E. (2002). Gene Expression Omnibus: NCBI gene expression and hybridization array data repository. Nucleic Acids Res 30, 207–210. doi: 10.1093/nar/30.1.207

Goh, W. W. B., Wang, W., and Wong, L. (2017). Why Batch Effects Matter in Omics Data, and How to Avoid Them. Trends in Biotechnology 35, 498–507. doi: 10.1016/j.tibtech.2017.02.012

Gomari, D. P., Schweickart, A., Cerchietti, L., Paietta, E., Fernandez, H., Al-Amin, H., et al. (2022). Variational autoencoders learn transferrable representations of metabolomics data. Commun Biol 5, 645. doi: 10.1038/s42003-022-03579-3

Hatano, M., Nagaoka, A., Miyahara, K., Hosogai, Y., Shishido, R., Hamasaki, H., et al. (2025). Impact of Confounding Factors in Human Postmortem Brain Tissues on Gene Expression Profiles: A Comparison of Patients With Schizophrenia, Bipolar Disorder, and Nonpsychiatric Controls. Neuropsychopharmacol Rep 45, e70053. doi: 10.1002/npr2.70053

Imani, M., Joudaki, M., Bagheri, A., and Arabnia, H. R. (2026). Why ROC-AUC Is Misleading for Highly Imbalanced Data: In-Depth Evaluation of MCC, F2-Score, H-Measure, and AUC-Based Metrics Across Diverse Classifiers. Technologies 14, 54. doi: 10.3390/technologies14010054

Jeni, L. A., Cohn, J. F., and De La Torre, F. (2013). Facing Imbalanced Data–Recommendations for the Use of Performance Metrics., in 2013 Humaine Association Conference on Affective Computing and Intelligent Interaction, 245–251. doi: 10.1109/ACII.2013.47

Khoroshevskyi, O., LeRoy, N., Reuter, V. P., and Sheffield, N. C. (2023). GEOfetch: a command-line tool for downloading data and standardized metadata from GEO and SRA. Bioinformatics 39, btad069. doi: 10.1093/bioinformatics/btad069

Leek, J. T., Scharpf, R. B., Bravo, H. C., Simcha, D., Langmead, B., Johnson, W. E., et al. (2010). Tackling the widespread and critical impact of batch effects in high-throughput data. Nat Rev Genet 11, https://doi.org/10.1038/nrg2825. doi: 10.1038/nrg2825

Marisa, L., de Reyniès, A., Duval, A., Selves, J., Gaub, M. P., Vescovo, L., et al. (2013). Gene expression classification of colon cancer into molecular subtypes: characterization, validation, and prognostic value. PLoS Med 10, e1001453. doi: 10.1371/journal.pmed.1001453

Papiez, A., Marczyk, M., Polanska, J., and Polanski, A. (2019). BatchI: Batch effect Identification in high-throughput screening data using a dynamic programming algorithm. Bioinformatics 35, 1885–1892. doi: 10.1093/bioinformatics/bty900

Parker, H. S., and Leek, J. T. (2012). The practical effect of batch on genomic prediction. Stat Appl Genet Mol Biol 11, Article-10. doi: 10.1515/1544-6115.1766

Smialowski, P., Frishman, D., and Kramer, S. (2010). Pitfalls of supervised feature selection. Bioinformatics 26, 440–443. doi: 10.1093/bioinformatics/btp621

Soneson, C., Gerster, S., and Delorenzi, M. (2014). Batch Effect Confounding Leads to Strong Bias in Performance Estimates Obtained by Cross-Validation. PLOS ONE 9, e100335. doi: 10.1371/journal.pone.0100335

Starcke, J., Spadafora, J., Spadafora, J., Spadafora, P., and Toma, M. (2025). The Effect of Data Leakage and Feature Selection on Machine Learning Performance for Early Parkinson’s Disease Detection. Bioengineering (Basel) 12, 845. doi: 10.3390/bioengineering12080845

Teschendorff, A. E., Journée, M., Absil, P. A., Sepulchre, R., and Caldas, C. (2007). Elucidating the altered transcriptional programs in breast cancer using independent component analysis. PLoS Comput Biol 3, e161. doi: 10.1371/journal.pcbi.0030161

Toubia, J., Conn, V. M., and Conn, S. J. (2018). Don’t go in circles: confounding factors in gene expression profiling. EMBO J 37, EMBJ201797945. doi: 10.15252/embj.201797945

Tsuyuzaki, K., Sato, H., Sato, K., and Nikaido, I. (2020). Benchmarking principal component analysis for large-scale single-cell RNA-sequencing. Genome Biol 21, 9. doi: 10.1186/s13059-019-1900-3

Yamamoto, H., Yamaji, H., Abe, Y., Harada, K., Waluyo, D., Fukusaki, E., et al. (2009). Dimensionality reduction for metabolome data using PCA, PLS, OPLS, and RFDA with differential penalties to latent variables. Chemometrics and Intelligent Laboratory Systems 98, 136–142. doi: 10.1016/j.chemolab.2009.05.006

Yu, Y., Mai, Y., Zheng, Y., and Shi, L. (2024). Assessing and mitigating batch effects in large-scale omics studies. Genome Biol 25, 254. doi: 10.1186/s13059-024-03401-9

